# Epigenetic heritability of cell plasticity drives cancer drug resistance through one-to-many genotype to phenotype mapping

**DOI:** 10.1101/2023.11.15.567140

**Authors:** Erica Oliveira, Salvatore Milite, Javier Fernandez-Mateos, George D Cresswell, Erika Yara-Romero, Georgios Vlachogiannis, Bingjie Chen, Chela James, Lucrezia Patruno, Gianluca Ascolani, Ahmet Acar, Timon Heide, Inma Spiteri, Alex Graudenzi, Giulio Caravagna, Andrea Bertotti, Trevor A Graham, Luca Magnani, Nicola Valeri, Andrea Sottoriva

**Author notes:** These authors contributed equally to this work (Erica Oliveira, Salvatore Milite, Javier Fernandez-Mateos).

## Abstract

Cancer drug resistance is multi-factorial, driven by heritable (epi)genetic changes but also phenotypic plasticity. Here we dissect it by perturbing colorectal cancer patient-derived organoids longitudinally with drugs in sequence. Combining longitudinal tracking, single cell ’omics, evolutionary modelling, and machine leaning, we found that different targeted drugs select for distinct subclones, supporting rationally designed drug sequences. The cellular memory was encoded as a heritable epigenetic configuration, from which multiple transcriptional programmes could run, supporting a one-to-many (epi)genotype-to-phenotype map that explains how clonal expansions and plasticity manifest together. This may ensure drug resistance subclones can exhibit distinct phenotypes in changing environments while still preserving the cellular memory encoding for their selective advantage. Chemotherapies resistance was instead entirely driven by plasticity. Inducing further chromosomal instability before drug application changed clonal evolution but not convergent transcriptional programmes. Collectively, our data show how genetic and epigenetic alterations are selected to “permissive epigenome” enabling phenotypic plasticity.

## Introduction

Cancers develop following the Darwinian rules of clonal evolution, where selected subclones bearing new heritable alterations come to dominate the cellular compartment^1^. The same paradigm is extended to explain the emergence of treatment resistance, arguably the biggest problem in oncology today^2^. However, only a subset of resistance mechanisms have been identified^3–5^. Even when genetic mutations are known to cause resistance, often these are only detected in a minor proportion of cells within tumours that are refractory to treatment. In many patients, treatment failure remains entirely unexplained by genetic alterations alone. Accumulating evidence suggests that additional Darwinian mechanisms involving non-genetic alterations^3,6,7^, as well as non-Darwinian cellular plasticity^8^ also contribute to tumourigenesis. Importantly, these mechanisms can co-exist in the same tumour at the same time and, hence therapy resistance is likely multi-factorial.

Relatively little attention has been paid to identifying epigenetic changes that drive cancer evolution, and “epigenetic driver” identification has been hindered by the lack of proper controls to compare cancer and normal epigenomes from the same tissue of origin^9^.

Further, because cancer cell plasticity is inherently a dynamic property, it is challenging to study in clinical samples typically collected at a single timepoint^10^. Elegant experiments based on the Luria-Delbruck approaches have demonstrated in cell lines that indeed one can distinguish plasticity from Darwinian adaptive changes^11^. Such mechanisms have also been identified in patient-derived xenograft models under the pressure of anti-EGFR drugs^12^.

Fundamental open questions remain, such as “to what extent are epigenetic changes heritable upon cell division?”, “is plasticity a new cancer programme or a reactivated cellular state?”, “is plasticity reversible?”, “is the propensity to cell plasticity in itself a heritable trait?”. Since therapy resistance is multi-factorial, another important question is: “are different subclones in the same tumour adapting differently to drugs?”. Answering these complex questions requires concomitant measurements of genomes, epigenomes and transcriptomes, matched with cell lineage histories to deconvolute Darwinian from non-Darwinian mechanisms. Because drug resistance is multi-factorial in the same tumour, single cell resolution is also required.

In this study we designed an evolutionary experiment using patient-derived tumour organoids under the pressure of different sequences of drugs. We used expressed lentiviral barcodes combined with single cell multi-omics to track the evolution of single cell genomes, epigenomes and transcriptomes over time. We studied adaptation in persistor cells under the pressure of drugs, and in the same cell lineages when drug pressure is released to understand how the therapeutic environment dynamically perturbs cells. We leveraged archetype analysis to chart the mapping between data modalities in individually tracked subclones.

## Results

### Experimental Design

Microsatellite stable (MSS) colorectal tumours are characterised by large numbers of chromosomal rearrangements, whereas microsatellite unstable (MSI) malignancies have largely diploid genomes but very high number of point mutations due to deficiency in the mismatch repair machinery. These are the two main genomic subtypes in colorectal cancer. In this study we utilise two patient-derived organoid lines. Firstly, an MSS, AKT-mutant and AKT amplified organoid that was derived from a metastatic colorectal cancer sample (patient F-016 in ref^13^). Secondly, an MSI mismatch repair deficient (MMRd) organoid that was derived from a primary colorectal cancer (CRC0282 from ref^12^). We infected each organoid with a library of 5 million random Cellecta® expressed barcodes, a form of lentiviral barcoding system that generates poly-A transcripts that can be captured by single cell RNA sequencing (see Material and Methods). We ensured to have one barcode per cell through dilution of 0.1 MOI. Single infected cell lineages are subject to high genetic drift due to cell turnover and death, giving rise to an expected final distribution of barcode sizes that scales as a power-law (Figure 1A). This power-law was used as the “null” distribution for comparison against our subsequent evolutionary experiments. We expanded each barcoded population to 75 million cells and 2.5 million cells/well were plated in multiple 6-well plates. At baseline, the “parental sample” of the first organoid line (AKT-mutant) had 3,000-5,000 individual cell lineages (Supplementary Figure 1), whereas the second organoid (MMRd) had lower barcode diversity (400 barcodes).

**Figure 1.**
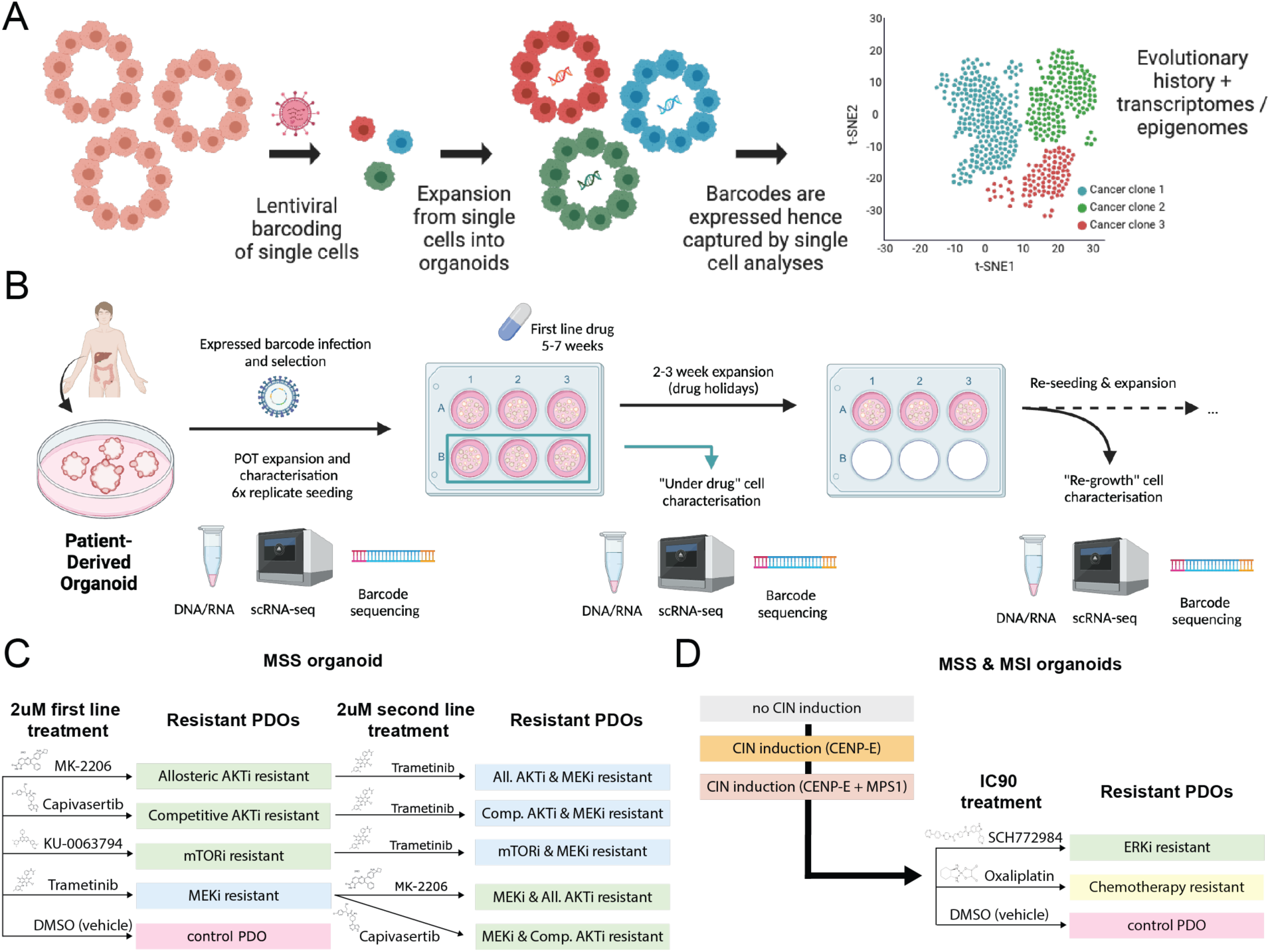
Experimental design of long-term drug resistance evolution in colorectal cancer organoids. **(A)** Lentiviral barcoding of single cells as an evolutionary tool. **(B)** Experimental design of a long-term drug resistance evolution in MSS AKT mutant organoid. We performed bulk DNA profiling for genomic characterisation and barcode measurement, as well as single-cell RNA-seq and corresponding single cell barcode extraction of five “solid” timepoints over a five-month period: parental, under drug 1, regrowth after drug 1, under drug 2 and regrowth after drug 2. We also collected floating DNA every two days from the supernatant to profile barcodes as a “liquid biopsy”. **(C)** We exposed the cells to 4 different sequences of drugs with first- and second-line treatments. **(D)** In a second experiment, we exposed both organoid lines (MSS and MSI) to an ERK inhibitor and oxaliplatin. Before drug pressure, we performed an induction of chromosomal instability (CIN) with CENP-E inhibitors and CENP-E + MPS1 inhibitors to study CIN effects on drug resistance.

We exposed the MSS AKT mutant organoid line to a set of targeted drugs for 45 days at a high dose of 2μM concentration (Figure 1B). We used allosteric AKT inhibitor (MK-2206), a competitive AKT inhibitor (capivasertib), an mTOR inhibitor (KU-0063794) and a MEK inhibitor (trametinib) (Figure 1C). We use those in sequence, making cells become resistant to drug A, followed by pressure by drug B (Figure 1B). Dose-response curves confirmed the cells became resistance to those drugs (Supplementary Figure 2A). We started with 6 replicas per condition, after 45 days collected cells under the pressure of the drug from 3 wells, and performed bulk DNA barcode analysis and single-cell RNA-seq. We left the other three wells regrowing in the absence of the drug until confluence and harvested the population again for barcode analysis, bulk DNA profiling and single-cell RNAseq. We re- plated cells into a new 6-well plate for a second drug and repeated the profiling as before. Every two days during the whole course of the experiment we collected “floating barcodes” by magnetic bead capture of cell-free DNA released in the media by apoptotic cells: this can be thought as a “liquid biopsy” of the cell culture. At the end of the experiment, we had five cellular timepoints directly derived from the system (i.e. a “solid” biopsy, Figure 1A) as well as 58-61 timepoints taken from floating barcodes. Floating barcode samples allowed characterising the system evolution without perturbing it with physical sampling.

In a subsequent experiment, both the MSS and MSI organoids were exposed to a single line of ERK inhibitors (SCH772984) and chemotherapy with oxaliplatin (Figure 1D) at IC90 concentration (Supplementary Table 1). This enabled the study of chemotherapeutic vs targeted drug. Furthermore, to probe the contribution of chromosomal changes to drug resistance, we induced chromosomal instability (CIN) in both organoid models by perturbing chromosome segregation^14^ using a CENP-E inhibitor alone or in combination with an MPS1 inhibitor^15^ (Figure 1D).

### Resistance evolution to targeted drugs but not chemotherapy is heritable, pre-existing and highly repeatable

We first focused on the MSS (AKT mutant) organoid. In all cases we found that drugs selected for one or a few subclones that massively expanded during treatment (Figure 2A). The three biological replicas we seeded per drug showed almost identical composition of selected barcodes, indicating that those subclones were pre-existing in the population and were selected by natural selection.

**Figure 2.**
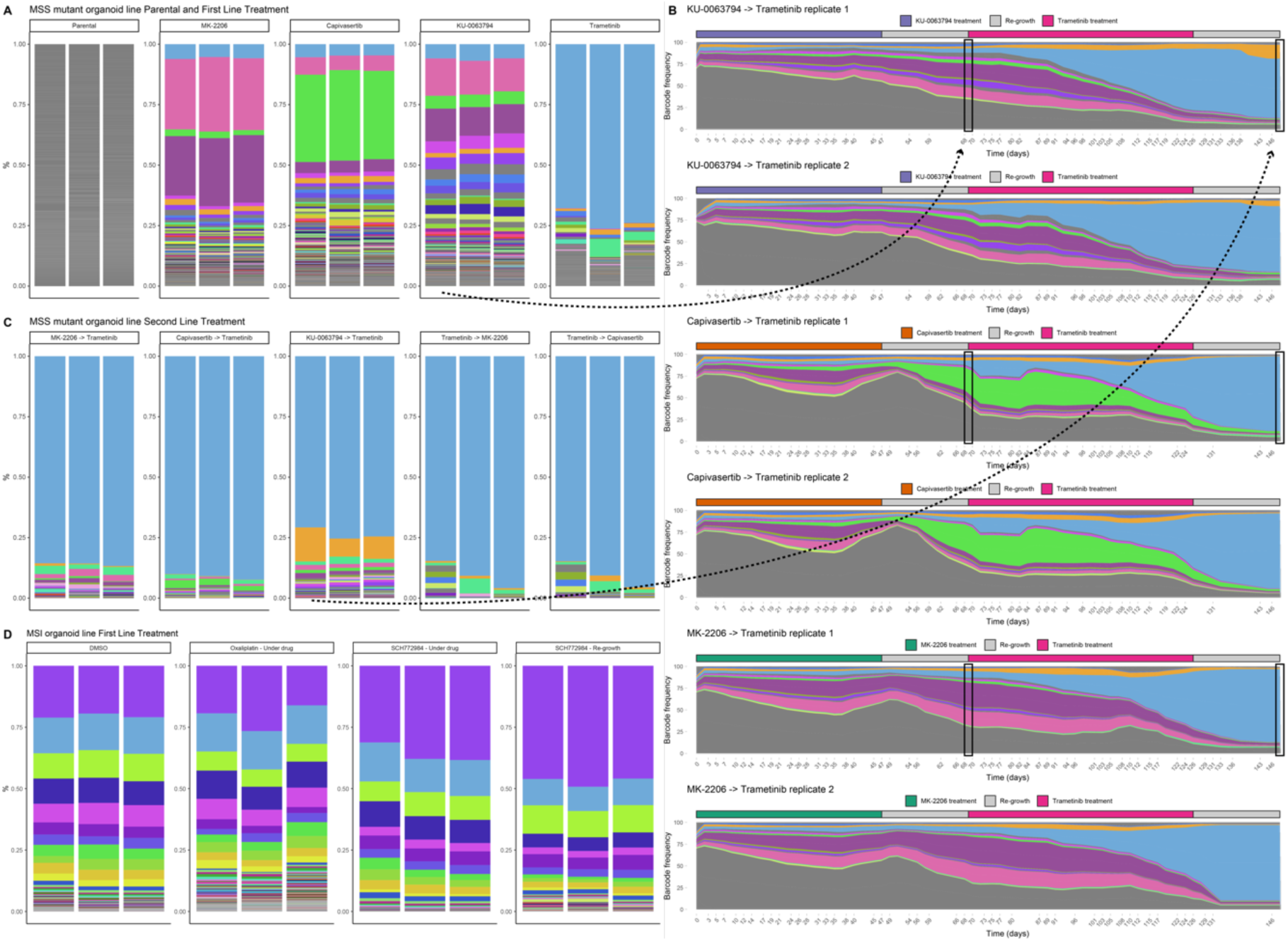
Evolutionary dynamics of barcoded population. **(A)** Lentiviral barcodes proportion following first line treatment in MSS AKT mutant organoid. For each replica and drug condition we quantified barcode proportions from genomic DNA, the only exception being trametinib for which we used the proportion quantified from 10x scRNA-seq. The top 100 barcodes have a unique colour across the whole experiment, all the others (< 2% abundance) are shown in grey. All the barcodes are quantified after the re-growth period. Selection is evident compared to the parental population. **(B)** Reconstruction of clonal dynamics using floating barcodes, extracted from culture media every 2 days over the whole length of the experiment. The dynamics shows an evident clonal sweep of the blue barcode after second line treatment with trametinib. Colour code is the same as in panel (A). Proportions are smoothed over a rolling average on a window of 7 points. **(C)** Lentiviral barcodes proportion with the second line treatment in MSS AKT mutant organoid. **(D)** Barcodes proportion in MSI organoid after being exposed to chemotherapy (oxaliplatin) and ERK inhibition (SCH772984).

We examined the floating barcodes to reveal longitudinal clonal dynamics. Dynamics were extremely similar between replicas, indicating that drug-resistance evolution was highly repeatable under the strong selective pressures of cancer drugs, and depended strongly on the initial conditions of the system (Figure 2B and Supplementary Figure 3). These data implied that the drug resistant phenotype was heritable, as the cellular memory encoding the drug resistant features was passed on to the offspring (i.e. drug resistance was clonally inherited).

Subsequent “second line” treatment with trametinib (MEK inhibitor) selected the same subclone as first line, which was cross-resistant to the other drugs when selected at first line (Figure 2C). We measured the growth rates over time of different clonal subpopulations (barcodes), showing that most surviving clones were not growing under the drug pressure, but were instead drug tolerant (i.e. barcode frequencies did not increase with time). When the drug pressure was released, the same subclones massively expanded with a high selective coefficient of up to *s=0.5* (Supplementary Figure 4). Similarly, in the MSI organoid, treatment with the ERK inhibitor SCH772984 revealed selection for pre-existing resistant subclones in a recurrent fashion between replicas (Figure 2D and Supplementary Figure 5). Contrastingly, chemotherapy with oxaliplatin did not select any pre-existing subclone in either of the two organoids, with the population structure of the cancer that remained unchanged, suggesting that instead plasticity could be responsible for resistance (Supplementary Figure 5).

### Different targeted drugs select for different pre-existing clones

We then investigated which subclones (identified by specific barcodes) were selected by different drugs. The two different AKT inhibitors, despite targeting the same signalling mechanism, selected for distinct subclones (Figure 2A). In contrast, the mTOR inhibitor selected similar subclones as the allosteric AKT inhibitor. The MEK inhibitor trametinib selected for an entirely different subclone that became dominant after drug exposure. The selective advantage of the trametinib-resistant subclone was very strong (Figure 2A and C; blue clone). The same subclone also proved to be a persistor in other drug contexts. For example, during first line treatment of AKT and mTOR inhibition, the trametinib-resistant subclone did not have a selective advantage but remained at low-frequency in the population until the second line of MEK inhibitor was later introduced which, subsequently, took over very rapidly.

These data indicate that evolutionary adaptation to targeted drugs was highly repeatable, driven by strong selective pressure on pre-existing subclones and led to the evolutionary divergence of subclones under distinct drugs. Therefore, evolution is determined by the initial conditions of the system (presence/absence of subclones), making it highly predictable, with the opportunity of exploiting evolutionary trade-offs for subclones in different drug contexts with specific sequences of drugs. These are the necessary conditions to exploit evolutionary *steering* strategies^16^.

We performed bulk whole-genome sequencing (WGS) of all MSS organoid samples at re- growth and compared to parental to identify candidate genetic alterations causing resistance. We found a small set of somatic mutations and copy number alterations that were different in resistant samples (Supplementary Figure 6 and 7). However, none of these variants were convincingly linked to a candidate mechanism of drug resistance.

### Archetype analysis reveals transcriptional programmes

In MSS organoids, we had, at first, performed single cell RNA-seq and single cell barcode extraction from a total of 19 samples (Supplementary Figure 8) and we were able to recover the barcode of 46% of cells on average per sample [30-64%]. Transcriptomes of cells under treatment clustered separately from parental, reflecting the strong influence of the environment on the transcriptomic profile of a cell (Figure 3A). Cells under different drugs also clustered separately, suggesting each drug influences the transcriptome differently (Figure 3B). Single cell barcodes recapitulated barcode distributions from bulk profiling (Figure 3C). As expected, most cells in S phase belonged to samples from re-growth after drug pressure (Figure 3D). We used markers of known cell types in the intestine to classify the transcriptional programmes active in each sample (Figure 3E). The parental sample was characterised by a small but significant component of LGR5+ stem-like cells and a large component of transient amplifying cells (TA). Under the pressure of first line AKT (capivasertib and MK-2206) and mTOR inhibitors (KU-0063794), which act on the PI3K/AKT pathway, the population maintained a stem cell component but expanded significantly a population with a Paneth signature, similar to what previously reported for patient-derived xenografts under the pressure of EGFR inhibitors^12^. Inhibiting the MAPK pathway with the MEK inhibitor (trametinib), even after PI3K/AKT suppression, induced instead different transcriptional dynamics where the LGR5+ stem-like component and partly a Goblet-like component emerged. Instead, blocking AKT after having previously inhibited MEK gave rise to yet another transcriptional programme with low LGR5+ stem component but high ASCL2+ stem-like cells. At regrowth, stem-like programmes were evident and a tendency of microfold cells. Instead for second line drugs a wider range of stem-like programmes emerge, again together with Paneth cell programmes.

**Figure 3.**
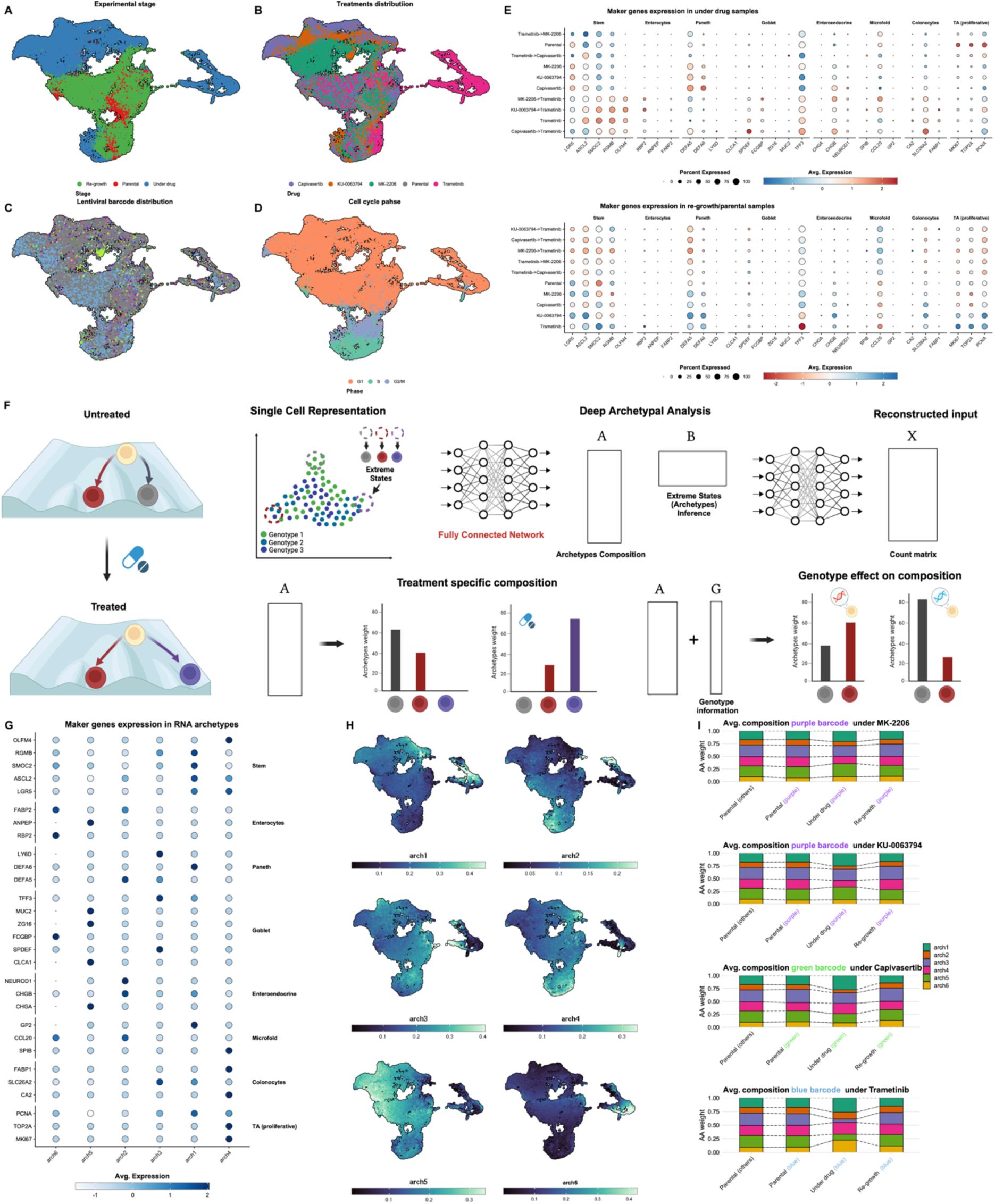
Transcriptional programmes show plasticity after drug administration. (A-D) UMAP of the 37,000 cells in the experiment after QC filters coloured respectively by: experimental stage, drug, barcode and cell cycle phase. Cells for which we were not able to extract a valid barcode or with an abundance of less than 1% are shown in grey. Cells in under drug phase tend to strongly cluster by drug, while they tend to mix back with the parental cells during re-growth. **(E)** Z-score distribution of adult colonic cell type markers from ref^20^ shows the presence of distinct differentiation programmes inside the organoid. **(F)** Archetypal Analysis (AA) aims at decomposing the input dataset as a convex combination of extreme points by learning two matrices A and B, which are the archetype weights for each point in the dataset and the matrix that defines the archetypes starting from the input dataset, respectively. Here we use a deep learning implementation of AA. We then exploited the weights of the A matrix to quantify differences in transcriptional programmes across conditions and genotypes **(G)** Z-score distribution for the same genes as in panel **(E)** but computed by archetype. **(H)** Archetypes weight distribution over UMAP. **(I)** Average archetype weight for different selected barcodes. The trend is consistent with cells going back to the parental phenotype after re-growth. Colours for barcodes are consistent with Figure 2.

Given the heterogeneity of transcriptional programmes we found, we hypothesised that a high level of phenotypic plasticity, possibly a reflection of aberrant differentiation pathways, was present in our cell populations. To deconvolve this signal we developed a new approach based on Archetypal Analysis, a dimensionality reduction technique that decomposes an input matrix of gene expression values as a convex combination of ideal extreme gene expression phenotypes called archetypes^17^. In this study we implemented a new Deep Archetypal Analysis framework called MIDAA^18^ to account for non-linearity in the dimensionality reduction step^19^ (Figure 3F). We found that distinct archetypes were enriched for clearly different cell type markers, highlighting the fact that they represent highly distinct cellular programmes, spanning all eight of the cell types we considered (Figure 3G). Archetype presence clustered distinctively between cells as expected (Figure 3H). These results indicate that archetypes are robust, cluster populations of cells in a meaningful way that is biologically interpretable. We performed the same analysis in the second line of experiments, obtaining similar results (Supplementary Figure 9). We used archetypes in the rest of the study as a surrogate of cell phenotype or transcriptional programme and studied their relationship to clonal evolution (with barcodes), copy number alterations and chromatin profiles.

### The drug environment produces plastic shifts in transcription that are reversible

We examined the composition of clonal barcodes within cells displaying each gene expression archetype. Subclones did not split by archetype, instead cells from the same barcoded clone gave rise to different archetypes (Figure 3I) suggesting “differentiation” to an archetype occurred in each cell lineage. Drug exposure (i.e. the environment) shifted the archetype distribution within a subclone, while those distributions were restored to the pre-treatment distribution following drug suspension and re-growth (Figure 3I). Hence, the selected subclones were transcriptionally plastic as in pre-treatment the archetype distribution within a barcode was similar to the background parental population, then changed under the drug, but finally returned to the original distribution of states after drug pressure was released. We observed a similar pattern in the second experiment where we exposed the two organoid lines to ERKi and chemotherapy, although this was expected under chemotherapy due to lack of selection for specific barcodes (Supplementary Figure 9E and 9F).

### The epigenome but not the transcriptome of selected subclones reveals their identity

We had observed that cells showed phenotype plasticity upon drug exposure, but drug resistance was nonetheless associated with clonal selection. We hypothesised that the epigenome could reconcile these apparently contradictory biological observations. We performed single cell multiome ATAC + Gene Expression on four samples: parental, trametinib, capivasertib and capivasertibètrametinib, as those were samples with the strongest bottlenecks. UMAP embeddings, in particular for the scATAC component, clearly separated the cells by treatment with a much stronger signal than the previous scRNA data (Figure 4A).

**Figure 4.**
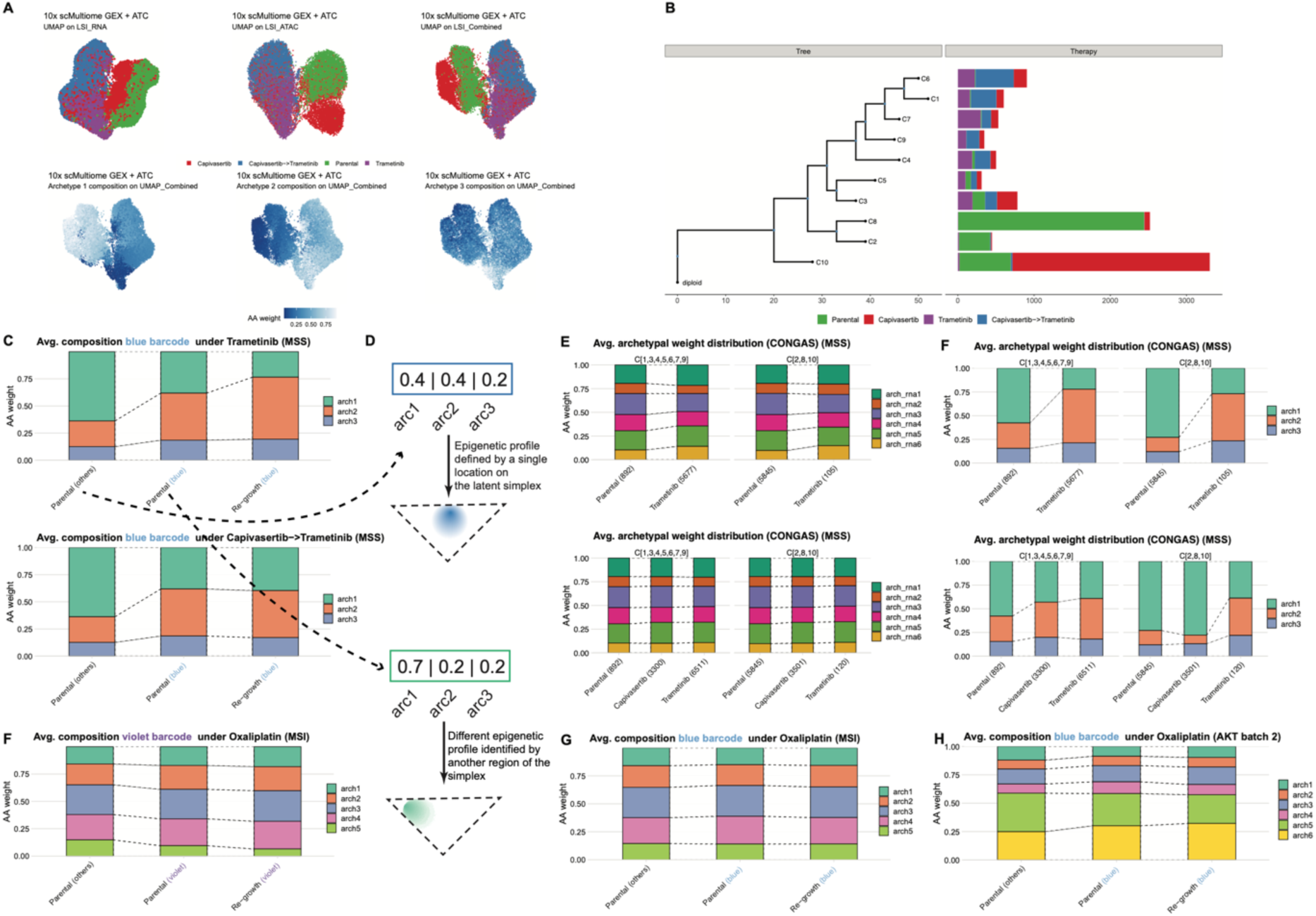
Epigenetic rewiring in resistant populations. **(A)** UMAP plots for multiome samples. In the first-row different dimensionality reductions are shown, respectively UMAP done with Latent Semantic Index (LSI) exploiting just RNA information, LSI with just ATAC and a combined LSI. In the second row we colour the combined UMAP by archetypal weight **(B)** Clonal tree constructed starting from CNAs inferred from ATAC data **(C)** Average archetype weights. The blue barcode displays a clear difference in the ATAC profile compared to the others (in grey). **(D-E)** Average ATAC archetype weight for copy number clones, for RNA archetypes **(D)** and ATAC archetypes **(E)**. We split the tree in two clades, the top one more abundant in the trametinib samples and the bottom one, more represented in the parental and capivasertib samples. The change in archetypal composition is consistent with what we saw with the lentiviral lineage tracing. **(F-G)** Average ATAC archetype weights for respectively the violet and blue barcode in the MSI sample **(H)** Average ATAC archetype weights for the violet barcode for the AKT organoid under oxaliplatin.

We inferred archetypes from the ATAC signal and called 3 archetypes that distributed differently in distinct samples. We then used the scATAC signal to infer copy number alterations with CONGAS^21^, identified 10 copy number alteration (CNA) subclones and reconstructed their phylogenetic history (Figure 4B). We used the lentiviral barcode recovery from the RNA component of the assay to assign barcodes to cells. The efficiency of barcode recovery here was lower (mean per sample = 27% [2-44%]) and we could only recover a strong signal for the ‘blue’ barcode selected under trametinib and capivasertib.

The proportion of epigenetic archetypes, unlike for scRNA, was different in the parental between the pre-existing selected barcode and the rest of the population, suggesting that the memory that induced the selection is encoded in the epigenome (Figure 4C). The epigenetic memory was maintained in the clone after it expands under AKT and MEK inhibition (Figure 4C). To perform the same analysis for other clones for which we did not have enough recovered barcodes, we performed subclonal decomposition of the cells using the copy number profiles and again compared transcriptional programmes using the RNA archetypes against their inherited CNA ‘genotype’ (Figure 4D). The transcriptional programmes were very similar when comparing selected versus non-selected subclones, whereas their epigenetic programmes were clearly different (Figure 4E), again indicating that there is a heritable epigenetic memory that encores many plastic transcriptional phenotypes downstream.

In the context of chemotherapy, both for the MSI organoid (Supplementary Figure 9E) and the MSS AKT mutant organoid (Supplementary Figure 9F), the plastic transcriptional rewiring was clearly evident, whereas both the barcodes (clonal structure of the population) and the chromatin remained stable throughout (Figure 4F-H), again confirming that plasticity – the potential to exhibit multiple phenotypes - was encoded by a single (epi)genomic configuration.

### Multiomic epigenetic analysis reveals heritable rewiring of the epigenome driving plastic phenotypes

We sought to define the epigenome configuration that enabled plasticity. First, we looked for recurrent focal changes in chromatin accessibility in promoters and enhancers in pre- vs post-treatment cells. We found a very small set of differentially accessible peaks in the promoters of differentially expressed genes (Supplementary Figure 10) of which only one was of particular interest: increased accessibility to the promoter of SETBP1, a known epigenetic regulatory hub^22^ (Supplementary Figure 11A). We then also looked at genome-wide rewiring of the epigenome by focussing on accessibility to transcription factor binding sites across the genome.

We found a significant correlation between gene expression of the TF and associated changes in the accessibility of the corresponding binding sites for a set of those TFs (Supplementary Figure 11B). Enriched TF accessible motifs are presented in Supplementary Figure 11C, and involve WNT and MAPK signalling pathways, as well as HomeoBox domains (Supplementary Figure 11D-E). Motif enrichment was then mapped into the UMAP showing localisation of chromatin accessibility patterns (Supplementary Figure 11F). A previous study in pancreatic cancer showed that resistance to trametinib was associated with increase in autophagy and decrease in MYC activity^23^. Similarly, we observed the autophagy expression signature under trametinib treatment (Supplementary Figure 11E). However, barcode analysis showed that the signal was not derived from the fully resistant subclone (the ‘blue’ trametinib resistant barcode), but from the residual persister cells that would eventually die after being overtaken by the ‘blue’ barcode (Supplementary Figure 11G,H).

### Induced chromosomal instability alters evolution but not phenotype

Chromosomal instability correlates with tumour progression^14,24^ and is likely to be involved in drug resistance as well. In recent work we showed that chromosomal configurations, although grossly altered, were relatively stable across long periods and through treatment in colorectal cancer patients^25^. Immune predation has also been shown to remove chromosomal aberrations^26^. The data suggested that negative selection was stabilising the karyotypes, and hence that cancers were sitting on a fitness peak, where all additional chromosomal variation was largely neutral, constrained around a fitness maximum. We had two questions: can we push cancers away from that maximum by inducing additional chromosomal instability? Would changing the initial chromosomal configuration in this way alter future evolution of drug resistance? We induced chromosomal instability with CENP-E and MPS1 inhibitors (see Material and Methods) in both organoid lines. The MSS AKT mutant organoid was MMRp and chromosomally highly altered to start with (Figure 5A), and exposure to the agents above proved relatively toxic (IC50=15nM for the MSS and 100nM for MSI organoid lines). In the MSS AKT mutant line this caused chromosomal rearrangements that nevertheless disappeared with time, and the cellular population returned to the chromosomal configuration of the parental line (Figure 5B). We found no alterations to the evolution of the tumour following exposure to drugs (Supplementary Figure 3). The MMRd organoid instead was chromosomally stable, mostly diploid (Figure 5C). It was much more tolerant to CIN-inducing agents and massive rearrangements in the chromosomal configuration were produced (Figure 5D) that had profound repercussions on future evolution. The population structure, as shown by barcode composition (Figure 5E-F) completely changed, and so did further evolution to targeted agents (Supplementary Figure 3). However, the transcriptional profiles driving resistance were not different than non-CIN induced samples (Figure 5G-H). This highlighted the different selective pressures acting on the cancer, and convergent evolution for drug resistance phenotypes which may be independent from DNA alterations.

**Figure 5.**
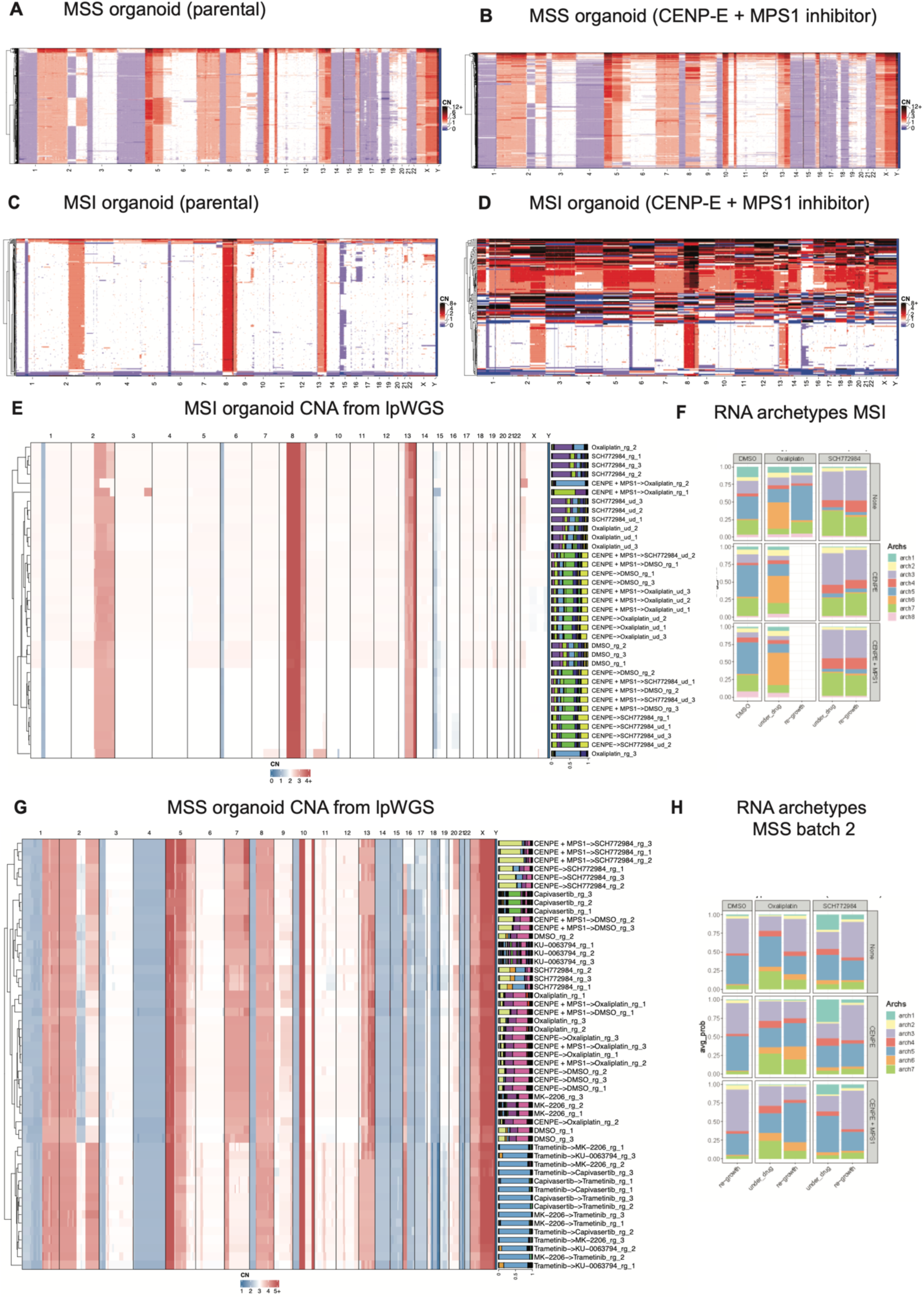
Inducing chromosomal instability before drug pressure. (A-B) Single-cell copy number profiles for the MSS AKT mutant organoid untreated and after treatment with CENP- E and MPS1 inhibitors. This organoid seems to be stable and resilient to drug induced alterations **(C-D)** Same as panel A and B but for the MSI organoid. In this case the situation is different and CENP-E and MPS1 inhibition causes a massive increase in instability with a residual bulk of population similar to the parental one. **(E-F)** Bulk copy-number profile and population structure as recapitulated by barcodes composition. We see how barcodes composition match copy-number clones, particularly evident with the blue trametinib resistant barcode in the MSS AKT mutant organoid. It is also clear how CENP-E inhibition induces significant changes in the population structure in the MSI organoid. **(G-H)** RNA archetypes distribution for MSI and MSS organoids. While as expected the drug induces a specific transcriptional phenotype, the induced instability acts only on the population structure, not influencing the transcriptome much.

### A one-to-many (epi)genotype to phenotype mapping drives resistance

Our evolutionary analysis demonstrates clear Darwinian dynamics driving resistance. At the same time the transcriptional programmes do not show such a pre-existing memory at the RNA level. Given the lack of genetic alterations responsible for resistance, we propose a model in which the combination of genetic alterations as SNVs and copy numbers, but importantly also the epigenetic configuration (Figure 6A) determine in what specific location on the fitness landscape a subclone resides (Figure 6B). Each subclone has a one-to-many map between the cellular memory encoded and heritable in its genome and epigenome, and the many possible transcriptional *phenotypes* it can produce in response to different environmental conditions. Therefore, the memory of phenotypic potential is retained long-term by a cell, but the exhibited phenotype displayed can change in response to environmental selective pressure. This potential for transcriptional heterogeneity enables rapid adaptation to changing microenvironments. Darwinian selection is for clone “memory”, namely the (epi)genomic configuration that enables a cell to access the phenotypic states that are adaptive in the face of drug selective pressure.

**Figure 6.**
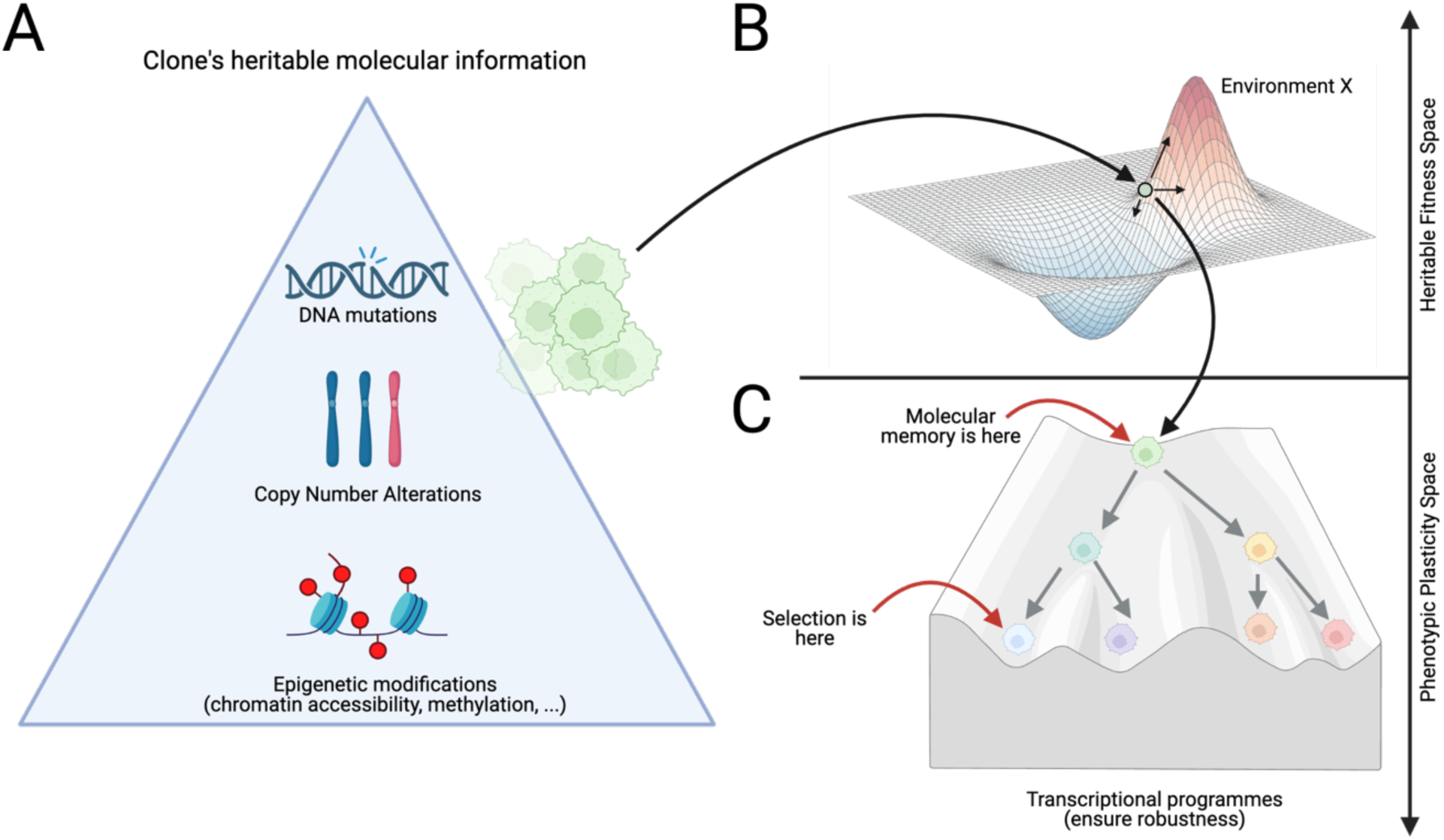
Heritability and plasticity of cellular phenotypes. **(A)** We propose a model in which genetic mutations and copy number alterations, together with heritable chromatin accessibility profiles determine the cellular memory of a certain clone, positioning it within a certain heritable fitness landscape **(B)**. However, the clone does not manifest as a single transcriptional phenotype, but rather as a set of transcriptional programmes that could be represented within a Waddington landscape, similarly to those that regulate development **(C)**. As Darwinian selection occurs at the level of phenotypes, it is likely that the selective pressure is at the bottom of the Waddington landscape, possibly for only a subset of the transcriptional programmes of a clone, despite the molecular memory carrying these programmes also containing as a side effect other plastic phenotype. This may explain the persistent phenotypic heterogeneity and plasticity of cancer clones, despite the strong selective pressure of treatments that instead should select for a single fittest phenotype.

## Discussion

Only a limited set of genetic alterations driving cancer drug resistance has been identified. Typical examples are gatekeeper mutations driving resistance to targeted therapies, such as in lung cancer^27^, or mutations in signalling pathways downstream of the targeted receptor, such as RAS/RAF mutations^2^, or mutations that revert a vulnerability^2^. For endocrine therapy in breast and prostate cancers, genetic alterations in ER^28^ and AR^29^ respectively have been described. However, in the majority of cases these mechanisms are not dominant (i.e. subclonal) in the tumour, supporting a view of polyclonal drug resistance^30^. Moreover, although chemotherapy is the backbone of cancer treatment in most malignancies, recurrent genetic alterations conferring resistance to chemotherapy have not been documented. This lack of insight is hindering the development of more effective treatments.

In this study we exploited patient-derived organoids, a new model system that is becoming established in the study of human diseases and in clinical research^13,31^, to design evolutionary experiments that are able to dissect the complex multi-faceted nature of drug resistance. This allowed us to discriminate between heritable genetic versus epigenetic evolution, and the interrelationship with phenotypic plasticity. We found that under the pressure of targeted drugs, the pre-existing heritable trait of being plastic is selected in a highly repeatable and predictable way.

The use of evolutionary theory applied to barcode information over time, combined with machine learning methods for single cell transcriptomics, shows the (epi)genotype-phenotype map is complex but readily explained by a genetic+epigenetic memory that is permissive for the expression of multiple phenotypes. Which phenotype is manifested by a cell depends on the environmental conditions, here the presence of cancer drugs. This finding could explain the high level of phenotypic plasticity present in tumours despite strong selection for individual resistance phenotypes, and at the same time the presence of clonal expansions that do not have a genetic driver explanation, and that could be driven by epigenetic heritable traits.

Overall, our study brings a mechanistic understanding of cellular plasticity within the clonal evolutionary framework^1,30,32^.

## Material and methods

### Organoid culture and passaging

MSS F-016 patient-derived organoid (PDO) was established from a liver metastasis of a colorectal tumour and it has been previously characterised as described in Vlachogiannis et al. 2018^13^. Among its pathogenic mutations, *AKT1* c.49G>A_p.Glu17Lys and *AKT1* amplification was of interest for our purpose as this organoid was a good model of oncogenic addition, showing high sensitivity and a selective apoptosis to Akt inhibitors^13^. CRC0282 patient-derived organoids was established from cohort in ref^12^.

PDOs were cultured embedded in Growth Factor Reduced (GFR) Basement Membrane Matrix (Corning), hereinafter referred to as matrigel, and Advanced DMEM/F12 media (Thermo Fisher Scientific), supplemented with 1X B27 and 1X N2 supplements (Thermo Fisher Scientific), 0.01% BSA (Roche), 2mM L-Glutamine (Thermo Fisher Scientific) and 100 units/ml penicillin streptomycin (Thermo Fisher Scientific). Additionally, 12 different growth factors were used to maintain PDOs culture: 50 ng/ml EGF, 100 ng/ml Noggin, 500 ng/ml R-Spondin 1, 10 ng/ml FGF-basic, 10 ng/ml FGF-10 (all from PeproTech), 10 nM Gastrin, 10 µM Y-27632, 4 mM Nicotinamide, 5 µM SB202190 (all from Sigma-Aldrich), 100 ng/ml Wnt-3A (R&D Systems), 1 µM Prostaglandin E2 and 0.5 µM A83-01 (Tocris Bioscience).

Passaging of PDOs was performed using TrypLE 1X diluted in 1mM PBS-EDTA (Thermo Fisher Scientific). In short, after media removal, PDOs in matrigel were harvested by pipetting with 1ml of TryplE1X and they were incubated for 20 min at 37°C, with mechanical homogenisation every 5 minutes. Then, PDOs were centrifuged at 1,200 rpm for 5 min at 4°C and washed with HBSS (Thermo Fisher Scientific). Counting and viability measurements were done using 0.4% Trypan Blue staining solution (Thermo Fisher Scientific) and the Countess 3 Automated Cell Counter (Thermo Fisher Scientific). Expected cells were pelleted again and re-seeded in matrigel.

### 3D patient-derived organoids drug screenings

To check baseline sensitivity to the drugs and inhibitors used, initial dose response curves (DRC) were performed with the allosteric inhibitor MK-2206, the non-allosteric one AZD5363 (capivasertib), ERK inhibitor SCH772984 and oxaliplatin . Initially, PDOs were dissociated into single cells as the passaging procedure described above, and after an automatic counting by Countess 3 Automated Cell Counter (Thermo Fisher Scientific, 6000 cells were seeded in 30µl of matrigel in 96-well plates. The matrigel was solidified after a 20-minute incubation at 37°C and 5% CO2, and overlaid with 70 µl of complete human organoid media. 24 hours later, media was removed and replaced with 50 µl of drug-containing media at different concentrations along with DMSO as a vehicle control, and replenished every two days for three times. Finally, drug-containing media was removed and replaced with 10% CellTiter-Blue cell viability assay media (Promega) and after 3-hour incubation at 37°C and 5% CO2, readings were taken in EnVision plate reader (PerkinElmer). Experiments were conducted in technical and biological replicates. DRC were represented by GraphPad Prism 9 (Dotmatics).

### Patient-derived organoids barcoding

Individual cell barcoding was performed by CloneTracker XP™ 5M Barcode-3’ Library in pScribe4M-RFP-Puro (Cellecta). This expressible barcode libraries enable the tracking and profiling of individual clones thanks to the integration in gDNA and barcode transcription in RNA. To ensure the insertion of a single barcode per cell, lentiviral titration was assessed and a MOI of 0.1 was chosen along with 0.8 μg/ml polybrene, corresponding to 10% of infection. 1 million cells from the organoids were infected following manufacturer’s protocol. After an over-night incubation in media suspension, cells were pelleted and resuspended in 1ml of matrigel in 6-well plates and with 2ml media containing 2.5μg/ml puromycin, as previously calculated by the puromycin killing curve. Following 10 days of puromycin selection, RFP-positive cells were checked under the microscope, and they were expanded with normal growth media. After several passages, parental population (POT) was frozen and characterised. DNA and RNA extractions and libraries sequencing (protocol below), detected around 3,500 different barcodes were in the final POT.

### Induction of chromosomal instability

F-016 (MSS) and CRC0282 (MSI) barcoded organoids were treated following Bennett et al. 2015 protocol. Briefly, following an overnight treatment with 100ng/mL of Nocodazole (487928, Sigma-Aldrich). Media was removed and replaced with fresh media containing IC50 concentrations (15nM for 3994-117 and 100nM for CRC0282) of CENP-E inhibitor GSK923295 (S7090, Selleckchem). After 2h, 150nM MPS1 inhibitor BOS172722 (S8911, Selleckchem) was added on top of one well for another 2h. Then, all the treatments were removed and replaced with fresh media. Organoids were allowed to grow for 10 days before performing the single cell sorting and freezing. The experiment was performed in 9 replicates and 3 replicates were collected at different timepoints along 15 days to check viability and barcodes distribution during the washout phase.

### Generation of drug resistant organoids

Once the F-016 POT was expanded, 75 million cells were used for the generation of resistant organoids to previously selected inhibitors. Briefly, 2,5 million cells/well were seeded in 6- well plates in 1ml of matrigel and 2ml of growth media. 24 hours later, media was collected and renewed with media containing 2μM of the selected inhibitors. For the first experiment, MK-2206 and capivasertib were chosen as Akt inhibitors, while trametinib, as a MEK inhibitor, and KU-0063794, as an upstream mTOR inhibitor, were also selected. DMSO was always used as the vehicle control.

Following above-described conditions, individual 6-well plates were used per each drug. Bottom three replicas were used for unique treatments while top three wells were used for a second treatment with drugs involved in different pathways, to perform a crossover screening. For the first experiment, F-016 PDOs were treated for 45 days with the respective first treatment and then, the three bottom wells were harvested and mixed to be used for bulk and single-cell characterisation, as well as for barcode amplification from DNA and RNA. Remaining cells were frozen and biobanked in FBS (Thermo Fisher Scientific), containing 10% DMSO (SigmaAldrich) for future analysis.

Top three wells were expanded without drug for two weeks (drug holidays), and then re- seeded with the same conditions as initial treatment, to be treated with the second inhibitor during additional 45 days. Lately, at the end of the treatment, cells were also expanded without drug for a couple of weeks and characterised and stored like initial time points.

As previously described, a total of 4 inhibitors were selected and a DMSO control plate was maintained until maximum confluency at day 15. For the crossover treatment, cells under Akt or mTOR inhibitors were treated with trametinib as a second treatment, while in cells initially treated with the MEK inhibitor, Akt or mTOR inhibitors were administered. To check the generation of resistant cells, sensitivity assays by dose response curves were performed at the end of each treatment.

For the second experiment, high doses corresponding to IC90 of oxaliplatin and the ERK inhibitor SCH772984 were used in both the MSS and MSI PDOs (Supplementary Table 1), with and without the CIN treatment previously explained, continually for 5 weeks. Then, the three bottom wells were harvested and used for bulk and single-cell characterisation, as well as for barcode amplification from DNA and RNA. Top three wells were expanded without drug for another 3 weeks before harvesting, characterisation and freezing.

### Parental and resistant cells characterisation by bulk and single-cell sequencing

Once treatments concluded, organoids were collected and dissociated into single-cell using the passaging procedure previously described. Half of the cells were used for bulk analysis while the other half was for single-cell experiments.

gDNA and RNA were isolated for bulk characterisation using All Prep DNA/RNA Mini Kit Qiagen (Qiagen) according to manufacturer’s recommendations. It mainly consisted of whole genome library preparation from 100ng of gDNA following NEBNext Ultra II FS (New England Biolabs) recommendations, with 20 minutes of enzymatic fragmentation and 5 PCR cycles. NEBNext® Multiplex Oligos for Illumina® (96 Unique Dual Index Primer Pairs, New England Biolabs) were used. After pooling, samples were sequenced for low-pass whole genome sequencing with at least 0.1X coverage in a NovaSeq 6000 (Illumina).

Regarding single-cell RNA approaches, a total of 19 single-cell experiments were performed from the parental (POT) and the 4 inhibitors (MK-2206, capivasertib, trametinib and KU- 63794) after drug removal (under drug) and after expansion (re-growth) for the two lines of treatment. For the second part, 36 single-cell experiments were performed, including the CIN generation and the following treatments with Oxaliplatin and ERK inhibitor SCH772984. After dissociation, single cells were washed with PBS and resuspended in PBS + 0.04% BSA, filtered through a 40µm FlowMi cell strainer (Sigma) and resuspended at a concentration of 1000 cells/μl. Viability was confirmed to be >90% in all samples using 0.4% Trypan Blue dye with Countess 3 Automated Cell Counter (ThermoFisher). Those organoids harvested under drug exposure with less than 70% viability were enriched by Dead Cell Removal Kit (Miltenyi) according to manufacturer’s protocol. For an estimation of 5000 cells, single cell suspensions were loaded on a Chromium Next GEM Chip G (10X Genomics) and were run in the Chromium Controller to generate single-cell gel bead-in-emulsions using the Chromium Next GEM Single Cell 3ʹ GEM, Library & Gel Bead Kit v3.1 kit (10X Genomics). After single-cell RNA-seq libraries were prepared and the library quality was confirmed with the TapeStation D1000 Screen Tape (Agilent) and a Qubit 3.0 dsDNA HS Assay Kit (Life Technologies), samples were pooled and sequenced on an Illumina NovaSeq 6000 (Illumina) according to standard 10X Genomics’ protocol.

Single-cell DNA approach was processed following 10x Genomics Single Cell CNV Solution kit (10X Genomics). Briefly, dissociated cells were partitioned in a microfluidic chip to form the cell beads and to continue with cell lysis and genomic DNA denaturation. Then, denatured gDNA in the cell bead followed a second encapsulation with barcoded gel beads to generate gel beads in emulsion (GEMs). Following a barcoding and amplification, fragments were later pooled to be linked to standard Illumina adaptors. Libraries were finally quantified by D1000 Screen Tape (Agilent) and a Qubit 3.0 dsDNA HS Assay Kit (Life Technologies), and pooled to be sequence on the NovaSeq S4 chemistry (Illumina) with 100 paired-end reads. Paired-end reads were processed using version 1.0 of the Cell Ranger DNA Pipeline (10× Genomics).

For single-cell multiome sequencing, after organoid dissociation, cells were lysed to obtain single nuclei following Chromium Nuclei Isolation Kit with RNase Inhibitor (10x Genomics, PN-1000494). Then, performing Chromium Next GEM Single Cell Multiome ATAC + Gene Expression (10x Genomics), around 5.000 nuclei followed a transposition step. Then, they were loaded on the Chip J Chromatin Controller (10x Genomics), for GEM generation and barcoding. After pre-amplification, ATAC and cDNA libraries were independently generated using user recommendations. Libraries were finally pooled and sequenced on the NovaSeq6000 (Illumina), following specific sequencing read cycles for an output of 25,000 read pairs per nucleus for both conditions.

### Barcode amplification and library preparation

Barcoded organoids at each time point before and after treatment were harvested and pelleted. To study the expressible barcodes, gDNA and RNA were isolated using All Prep DNA/RNA Mini Kit Qiagen (Qiagen) according to manufacturer’s recommendations. While DNA was directly quantified by Qubit 3.0 dsDNA HS Assay Kit (Life Technologies) and stored for barcode amplification, 300ng of total RNA were taken for cDNA synthesis with SuperScript^TM^ II Reverse Transcriptase (Invitrogen).

Contrary to a 2D culture, organoids are embedded in matrigel and although dead cells remained inside, gDNA is released to the surrounded media. Hence, to track the evolution of each cell lineage during the treatment without replating them, gDNA from apoptotic cells was collected every media change (2 days) to amplify dead cell barcodes, providing us unparalleled temporal resolution. Collected media was incubated for 2 hours at 55°C with 20mg/ml proteinase K (Roche). Then, gDNA was purified with 3X SPRIselect beads (Beckman Coulter) followed by an 80% ethanol wash step. DNA was eluted in low TE and quantified by Qubit 3.0 dsDNA HS Assay Kit (Life Technologies).

Barcode amplification by PCR was performed using 2x Accuzyme mix (Bioline) and 10 ng of the extracted DNA/cDNA using the primers below (10μM):

Forward XP primer: 5’-ACCGAACGCAACGCACGCA-3’

Reverse XP primer: 5’-ACGACCACGACCGACCCGAACCACGA-3’

Briefly, 25μl reactions PCR steps were: 98°C 2’, and 35 cycles of 95°C 15’’, 71°C 15’’, 72°C 10’’ and a final extension of 72°C 3’. 4μl of the 128-bp PCR product were later checked on an 1.5% agarose gel and quantified for library generation by TapeStation (Agilent). NGS libraries were prepared from 30ng of the PCR product using the NEBnext Ultra II DNA library preparation kit for Illumina (New England Biolabs) according to manufacturer’s recommendations and using NEBNext® Multiplex Oligos for Illumina® (96 Unique Dual Index Primer Pairs, New England Biolabs). Libraries were quantified using Qubit 3.0 dsDNA HS Assay Kit (Life Technologies) and TapeStation D1000 Screen Tape (Agilent Genomics). Up to 384 samples were pooled for sequencing. Due to unspecific library amplification, 3ng/μl samples were pooled considering 250-340bp concentration in TapeStation. Then, pools were dried out to 50ul using a vacuum concentrator and an electrophoresis in 2.5% gel was done. 280pb band was cut and purified by Gel purification kit (Qiagen). 50μl of the eluted product was again quantified by both methods. NGS was performed at the Tumour Profiling Unit of the Institute of Cancer Research using NovaSeq 6000 (Illumina). 150 pair-end reads and 15% of PhiX was considered and approximately 5000 reads per time-point was aimed.

### Phenotypic correlation with barcodes

CloneTracker XP™ barcode enrichment was performed in single-cell gene expression libraries. The 10x Chromium 3’ Reagent Kit was developed for amplification of 3’-ends of polyA+ RNAs. Due to the CloneTracker XP BC14-spacer-BC30 barcode design, located at a fixed ∼150-200nt distance upstream from the poly-A+ site, the transcribed barcodes are captured in the first step of the 10x Chromium 3’ protocol.

To effectively read the barcode sequence an additional PCR amplification step was needed with a barcode-specific primer that is located just upstream of the CloneTracker XP barcode (FBP1). This ensured amplification of the segment of cDNA that contains both the CloneTracker XP barcode and the cell barcode associated with the poly-A+ sequence in the scRNA-Seq protocol. In short, after cDNA generation by 10x Chromium Next GEM Single Cell 3ʹ protocol, 75% of the adaptor ligated product was used for gene expression libraries, while 25% of amplified cDNA was taken for sequential nested PCR barcode amplification. First PCR was performed for 9 cycles using below primers:

Partial 10X Read 1 primer forward: 5’- ACACTCTTTCCCTACACGACGCTCTTCCGATCT- 3’ Specific barcode fragment primer (FBP1) FSeqRNA-BC14-XP Reverse Primer:

5’- GTGACTGGAGTTCAGACGTGTGCTCTTCCGATCTCCGACCACCGAACGCAACGCACGCA- 3’

Second round of PCR to increase Clone Tracker XP barcoded cDNA sequences used standard 10X p7 and p5 primers for additional 6 cycles to then proceed to indexing and final library generation. After QC by TapeStation (Agilent), barcode amplicon libraries were pooled and sequenced. Then, Clone Tracker XP barcodes and gene expression were correlated thanks to 10X barcodes identifying cellular gene expression with the clone identifiers.

### Barcode bioinformatics analysis

Barcodes from bulk experiments were quantified using *BWA*^1^ and *FeatureCounts* ^2^. We first generated a fasta reference using the full pool of barcodes provided by Cellecta. We then aligned the amplicons using *bwa-mem* with custom parameters. The bam files obtained from bwa were then used to quantify barcode abundances using the *FeatureCounts* implementation of the R package *Rsubread* ^3^. More specifically we run the command *featureCounts* we filtered reads with mapping quality less than 30 (minMQS = 30).

For barcodes enriched from the 10x Chromium library we used STAR solo ^4^ with a custom reference index built on the reference fasta used for BWA.

### Barcode evolutionary modelling

We used the floating barcodes abundances to produce an estim1ate of the fitness value of each barcode. To avoid biases due to the differential efficacy in capturing the DNA across different phases of the experiment we worked with the relative abundances of each barcode and to make the calculations tractable for thousands of barcodes we assumed exponential growth. Under this assumption if call as ω_*i*_ the effective growth rate (so the difference between birth rate and death rate) for barcode i we can write its abundance at time T as:

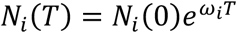

As we were working with relative abundances it follows, if we assume independence between subclones, that the abundance *f*_*i*(*T*) for barcode I at time T is simply:

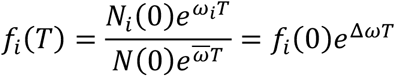

Where N(0) is the population size at time 0, ω is the average growth rate of the population and Δω id the difference between the barcoded population growth rate and the average population. From this simple model we derive that the relation between the abundance at sampling time T and time T-1 is (after taking the logarithm of both sides):

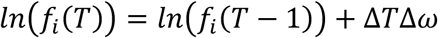

We then calculate the relative growth rate by fitting the beta coefficients of a linear model on the log abundances using the *fastLM* function of the *RcppEigen*^33^ R package.

### Single cell RNA sequencing analysis

For the analysis of single-cell RNA sequencing (scRNA-seq) data, the fastq files containing the sequential reads were aligned to the reference genome (assembly hg38) using STARsolo version 2.7.9a.

The matrices produced from the alignment were corrected for ambient RNA using the *adjustCounts* function of *SoupX*^34^. We then used *DoubletFinder*^35^ to estimate the doublet status of each cell. For quality control, filters were applied for the minimum number of genes with non-zero expression and for the percentage of expression derived from mitochondrial transcripts (%mt), indicative of dying cells. Specifically, cells with %mt greater than 30%, fewer than 1000 expressed genes and putative doublet from *DoubletFinder* were filtered out.

Data was then analyzed in Seurat^36^. Counts were normalized using the *NormalizeData* function. At this point, the 3000 most variable genes were identified with the FindVariableFeatures routine, which were then scaled (*ScaleData*) and used to compute a Principal Component Analysis (*RunPCA*). The first 25 components, selected based on the amount of variance explained, were used to compute the KNN graph and Uniform Manifold Approximation (UMAP) with the functions *FindNeighbors* and *RunUMAP*.

We compute gene set scores for the Autophagy signature from Ref.^23^ and the *MSigDB*^37^

Hallmark of Cancer MYC targets V2 using the function *AddModuleScore*.

### Single cell ATAC + GEX analysis

10X multiomics sequencing data has been processed with cellranger-arc version . We then performed all the quality control and analysis using the R package *ArchR*^38^ . After generating the arrow files, we used the *filterDoublets* and filtered out cells with *TSSEnrichment* < 6, number of fragments < 10000, putative doublets and cells without a valid UMI for both GEX and ATAC libraries. LSI for RNA and ATAC were computed using the *addIterativeLSI* method using respectively the 2500 most variable genes and the ArchR tile matrix. Motif annotations for *ChromVAR*^39^ were added using the *addMotifAnnotations* function with using the *CIS-BP* dataset and scores calculated using the *addDeviationsMatrix* method. To compute correlation, we used the *correlateMatrices* function with input the Motif Matrix and the Gene Expression Matrix. Marker peaks were computed using the *getMarkerFeatures* and setting as bias terms the TSS enrichment score and the log10(number of fragments). To then computer differential Transcription Factor Motif accessibility we used the function *peakAnnoEnrichment*.

Background peaks were computed using the *addBgdPeaks* method.

### Deep Archetypal Analysis

Archetypal Analysis is a dimensionality reduction method that decomposes an input matrix as a convex combination of ideal extreme points called archetypes and it was first introduced in ref^17^. More in detail if we assume *X* ∈ *R* to be our *N x M* input matrix, the classical problem of archetypal analysis revolves on optimizing the following loss function:

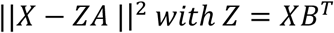

Where B and A are respectively *N x K* and *K x N* row stochastic matrices with K being the number of archetypes. We define a matrix X to be row stochastic matrix if it satisfies x_!&_ ≥ 0, ∑_&_ x_*ik*_ = 1. The matrix B defines the archetype as a convex combination of real data points, while the matrix A represents the weights of each archetype in reconstructing each cell. This is intuitively equivalent to optimally embed the data in a convex polytope with K vertices (archetypes), it then follows that the archetypes define the convex hull of the polytope.

The original algorithm for finding the minima involves alternating optimization steps were we first fix B and optimize for A and then do the do the reverse. This solution is generally considered hard to scale as it needs to use all the whole dataset for each iteration ref^40^.

To add the possibility of mini batch learning and to account possible non-linear effects in the archetypal composition we implemented the Deep Archetypal Network presented in ref^19^ into a new tool called MIDAA^18^.

In this case we learn both matrices using a feed-forward neural network *A,B* = *f*_θ_(*X*). The input is then reconstructed by a specular decoder network *f*_ϕ_(*Z*^∗^*A*) = *X*^F^. To reduce the complexity of the problem Z* is fixed and set to a standard simplex with K vertices and the condition *Z* = *XB*^*T*^ is guaranteed by using an additional term in the loss, that becomes:

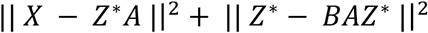

To fit the scRNA-seq data we first computed the intersection top 2000 most variable genes in the under-drug phase, the 1000 most variable in re-growth and the 500 most variable in the parental sample. We then computed the Z-score across all the cells with a valid barcode and used it as an input for the AA net. For the scATAC-seq from the 10x multiome, given the higher complexity we used the first 30 components of the LSI as input.

### Copy Number Analysis

#### Copy Number Analysis from ATAC Data

To infer copy number clones from the ATAC portion of the single-cell multiome we used CONGAS+^21^. To derive a prior segmentation, we used the inferred copy number states from the corresponding low-pass DNA analysis, and we did an intersection of the segments among the samples to obtain a unique profile. We then used the function *segments_selector_congas* to filter for putative multimodal segments. We run the method *fit_congas* and tested a range of clusters going from 1 to 15 for 2000 steps, a learning rate of 0.005 and a temperature parameter for the Gumbel-SoftMax of 10. We choose the best number of cluster that minimized the Integrated Completed Likelihood (ICL). The inferred copy number alteration matrix was used as an input to MEDICC2^41^ with parameters *--total-copy-numbers --input-allele-columns* to build a clonal tree.

#### Copy Number Analysis from 10x Single Cell DNA Data

As per the 10x recommended analysis, CNAs were determined using CellRanger with default settings (using GRCh38 as the reference genome). For each sample a threshold of maximum number segments per cell was used, as determined by manually assessing 30 random cells from each sample for noisy profiles. The thresholds were 190 for the MSI parental and CENPE-treated organoids, 299 for the CENPE- and MPS1-treated MSI organoid, 291 for the AKT parental organoid, 249 for the CENPE-treated AKT organoid and 227 for the CENPE- and MPS1-treated organoid. Calls were then binned into 1Mbp bins and bins with no overlapping segments in at least one cell were removed. Cells with more than 50% of the genome with copy number zero were removed, as were cells with a mean copy number of more than 20.

#### Copy Number Analysis from Low-Pass Whole-Genome Sequencing

FASTQ files were trimmed for adaptor content using skewer^42^ with a minimum length allowed after trimming of 35 bp, keeping only reads with a minimum mean quality of 10 and removing highly degenerative reads (-l 35 -Q 10 -n). Trimmed reads were aligned to hg38 (GRCh38) using bwa mem^43^. SAM files were sorted, compressed to BAM files and duplicates were marked using Picard tools (https://broadinstitute.github.io/picard/). BAM files were then indexed also using samtools^44^.

BAM files were processed using QDNAseq^45^ to convert read counts in 500kb bins across the chromosomes of hg38 into log2 ratio data. The 500kb bins for hg38 were generated according to QDNAseq instructions using the GEM library^46^ and normal BAM files from the 1000 genomes project (1000genomes.ebi.ac.uk, phase 3). Data normalisation was performed in accordance with the QDNAseq workflow, including sex chromosomes. Bins were required to have a minimum mappability of 65 and 95% non-N bases. The *smoothOutlierBins* function step was removed as it artificially depressed highly amplified bins. The *sqrt* option was used for the *segmentBins* function. Log2 ratios in bins and segments were normalised by subtracting the median log2 ratio value of all autosome bins.

To call absolute copy number we used an adapted version of the ASCAT^47^ approach using only log2 ratio information and calculating the sum of squared differences between per bin segmented copy number and the nearest positive integer within a grid search of purity and ploidy parameters and identifying the best fitting local minima. Based on expected ploidy statuses, we search for fits in the range of 2.9 – 3.3 for the AKT organoid samples and 1.9 –2.3 for the MSI organoid samples, with a minimum purity of 0.98 as these are pure samples. If no local minimum was found a ploidy of 3.1 for was used for AKT and 2.1 for MSI, with purity being set at 1 for both. Purity and ploidy searches were performed only on the autosome segmented bins and the X chromosome copy number status was inferred from the resulting solution. Continuous copy number values were reported to capture heterogeneity in Figure 5E and F.

For CONGAS input (AKT organoid only) we required segmentation and calling that was comparable across the multiple samples, therefore we employed multi-sample segmentation using the copynumber package (*multipcf*, gamma = 10) and called copy number as previously described but across a narrower ploidy range of 3.1±0.1.

## Acknowledgments

This work was supported by the AIRC/CRUK Accelerator Award (A26815). A.S. and T.G. are also supported by the Wellcome Trust (202778/B/16/Z and 202778/Z/16/Z respectively) and Cancer Research UK (A22909 to A.S., A19771 & DRCNPG-May21_100001 to T.G.). NV was supported by Cancer Research UK (A18052 and A26815), the National Institute for Health Research (NIHR) Biomedical Research Centre at The Royal Marsden NHS Foundation Trust and The Institute of Cancer Research, the European Union FP7 (CIG 334261) and the Katherine and Douglas Longden Chair in Oncology at Imperial College London. This work was conducted with funding from AIRC, Associazione Italiana per la Ricerca sul Cancro, AIRC 5x1000 grant 21091 (to A.Be.). GC acknowledges support from the Italian Association for Cancer Research (AIRC) - MFAG 2020 [grant number 24913].

## Conflict of Interest

Between 2018 and 2022 NV received honoraria for lectures from Merck Serono, Pfizer, Bayer, Eli-Lilly and Servier; Research funding (Institutional) from Roche and BenevolentAI and has received paid consultancy fees from BenevolentAI. At the time of submission NV was a full-time employee of AstraZeneca Plc; any work done in relation to this article was undertaken before his current employment.

## Code Availability

The code to reproduce the analysis and the figures is available on GitHub (https://github.com/sottorivalab/epigenetic_heritability_and_cell_plasticity_reproducibility). We also provide annotated Seurat objects and barcode tables in Zenodo (10.5281/zenodo.11058999)

## Supplementary Figures

**Supplementary Figure 1. *Distribution of lentiviral barcodes from the POT across biological and technical replicates.*** The plot shows how lentiviral barcode proportions in absence of selective pressure follow and expected power-law distribution and how the number of barcodes retrieved is consistent across replicates.

**Supplementary Figure 2. Drug response curves of the resistant organoids. (A)** Drug response curves related to the first experiment using the MSS AKT organoids after the long-term treatments with the 4 targeted inhibitors (MK-2206, capivasertib, trametinib, and KU-0063794) during the recovering stage showing they remained resistant after the drug removal in all replicates. (B) Drug response curves related to the second experiment after the CIN treatment and long-term exposure to oxaliplatin and SCH772984 after the 3 weeks of recovery without the drug, showing that the cells recovered sensitivity after the heavy treatment.

**Supplementary Figure 3. *Floating barcodes evolution for the whole cohort***. Fishplots as in Figure 2 panel B showing the clonal dynamics of all the replicates and the sample in the experiment.

**Supplementary Figure 4. *Relative fitness distribution computed over floating barcodes abundance.*** Relative fitness distribution shows high variability in re-growth phase during first-line treatment. Coefficients were calculated by assuming exponential growth and independence between different clonal populations. **(A)** MSS AKT organoid first batch, **(B)** MSS AKT organoid second batch, **(C)** MSI organoid.

**Supplementary Figure 5.** Lentiviral barcodes proportion for the AKT organoid after the CENP- E/MPS1 perturbation experiment and during the resistant generation with oxaliplatin and SCH772984. Colours are consistent with Figure 2 panel A and B, so that grey barcodes are lowly abundant barcodes are in fact defined compared to the first drug experiment.

**Supplementary Figure 6. *Allele frequency distribution in parental vs capivasertib->trametinib treated organoids.*** Scatterplot of the variant allele frequency of putative functional mutations. Mutations were selected as either having VEP MODERATE impact and being either deleterious according to SIFT or damaging according to PolyPhen (left panel) or having VEP HIGH impact (right panel). No obvious resistance gene is present off diagonal.

**Supplementary Figure 7. *Copy number profiles of treated and untreated organoids from low-pass WGS.*** Heatmap of relative copy number profiles for all the organoids in the experiment. Gain and losses are expressed compared to the Parental. The similarity across replicas and drug treatments is an orthogonal validation of the results obtained by lentiviral barcoding.

**Supplementary Figure 8. *Lentiviral barcodes distribution in 10X scRNA-seq.*** (A-B) Barcode distribution in the parental and the treated organoids respectively, the top 100 barcodes in frequency are colored the others are shown in grey. (C) UMAP colored by barcode as in Figure 3C. Colors are consistent with Figures 2-3 and across panel, such that each color is always a unique barcode.

**Supplementary Figure 9. Transcriptional Landscape of organoids treated with CENP-E and MPS1 inhibitor. (A-B)** Z-score distribution of markers in Figure 3E, we note how the MSI organoid seems to be less heterogenous. **(C-D)** Cell type specific marker expression for each archetype in the MSI and AKT (second batch) organoid, in both cases we see how archetypes capture some cell-type specific variability. **(E-F)** Archetype distribution for selected barcodes over the course of the experiment in MSI **(E)** and AKT batch 2 **(F)** organoids. We find again a massive transcriptional rewiring after therapy that tends to slowly come back to an untreated like status after re-growth. It is important to note that this effect is not homogenous across drugs and organoids in this case with the AKT rapidly coming back to a DMSO like state after SCH77298, while in the MSI case it seems like the drug the is the induces this strong on-off transcriptional requiring is the Oxaliplatin.

**Supplementary Figure 10. *Trackplot of peaks enriched in gene promoters after Trametinib treatment.*** We first run differential expression analysis on the Multiome GEX part using a non-parametric Wilcoxon test grouping by drug. We then perform the same analysis but with peak coverage. In both cases we selected the results with FDR <= 0.01 and absolute logFC > 1. To generate the final set we intersected the significant peaks and genes, by considering only peaks in the promoter regions.

**Supplementary Figure 11. Biological characterization of Trametinib resistant population. (A)** *SETBP1* chromatin profile in the promoter region. **(B)** Scatterplot of Transcription Factor (TF) Motifs variance computed using ChromVAR^39^ from the ATAC portion of the Multiome and expression from the GEX part. Points in red are enriched TFs that also show consistent high change in expression, selected as having correlation > 0.6, adjusted p-value < 0.001 and TFs Motif variance greater than the 10% quantile. **(C)** Heatmap of enriched Motifs in Marker Peaks. We filtered Enriched Motifs with FDR <= 0.001 & Log2FC >= 1 **(D-E)** StringDB analysis of Motifs enriched respectively in capivasertib->trametinib and trametinib samples, showing involvement of TFs active in WNT and MAPK signalling pathways as well as TFs containing HomeoBox domains. Analysis was conducted using the browser version 12.0 of StringDB^48^ **(F)** UMAP plots for some of the top enriched Motifs. Sample specific localization is evident. **(G-H)** Gene Module score for the autophagy signature in Ref.^23^ and the MSigDB Hallmark of Cancer MYC targets V2^37^, grouped respectively by phase and barcode. The plot shows how in the under-drug phase cells under Trametinib have and high level of autophagy and a correspondingly lower level of MYC activation. The blue resistant barcode however shows an opposite trend and seems to resemble the parental and re-growth behavior even under drug. Panel **(H)** scores are computed for the under-drug samples.

**Supplementary Table 1.** IC90 values assessed by CellTiter-Blue cell viability assay. Organoids were treated with serial dilutions of the drugs along with DMSO as a vehicle control, and treatment was replenished every two days for three times. Experiments were conducted in technical and biological replicates. Plates were assessed by CellTiter-Blue cell viability assay media (Promega).

| IC<sub>90</sub> values | Oxaliplatin (μM) | SCH772984 (μM) |
| --- | --- | --- |
| MSI Parental | 25 | 0.5 |
| MSI CENPEi | 25 | 1 |
| MSI CENPEi + MPS1i | 50 | 1 |
| AKT Parental | 50 | 1 |
| AKT CENPEi | 50 | 1 |
| AKT CENPEi + MPS1i | 50 | 1 |

## Notes

### Summary of Updates

Additional results were added (see new Figure 5) and methods were extended / clarified.

